# Estimating directed connectivity from cortical recordings and reconstructed sources

**DOI:** 10.1101/023523

**Authors:** Margarita Papadopoulou, Karl Friston, Daniele Marinazzo

## Abstract

In cognitive neuroscience, electrical brain activity is most commonly recorded at the scalp. In order to infer the contributions and connectivity of underlying neuronal sources within the brain, it is necessary to reconstruct sensor data at the source level. Several approaches to this reconstruction have been developed, thereby solving the so-called implicit inverse problem (Michel et al. 2004). However, a unifying premise against which to validate these source reconstructions is seldom available. The dataset provided in this work, in which brain activity is simultaneously recorded on the scalp (non-invasively) by electroencephalography (EEG) and on the cortex (invasively) by electrocorticography (ECoG), can be of a great help in this direction. These multimodal recordings were obtained from a macaque monkey under wakefulness and sedation. Our primary goal was to establish the connectivity architecture between two sources of interest (frontal and parietal), and to assess how their coupling changes over the conditions. We chose these sources because previous studies have shown that the connections between them are modified by anaesthesia (Boly et al. 2012). Our secondary goal was to evaluate the consistency of the connectivity results when analyzing sources recorded from invasive data (128 implanted ECoG sources) and source activity reconstructed from scalp recordings (19 EEG sensors) at the same locations as the ECoG sources. We conclude that the directed connectivity in the frequency domain between cortical sources reconstructed from scalp EEG is qualitatively similar to the connectivity inferred directly from cortical recordings, using both data-driven (directed transfer function; DTF) and biologically grounded (dynamic causal modelling; DCM) methods. Furthermore, the connectivity changes identified were consistent with previous findings (Boly et al. 2012). Our findings suggest that inferences about directed connectivity based upon non-invasive electrophysiological data have construct validity in relation to invasive recordings.

## Introduction

Oscillatory synchronous activity of local or distributed neuronal populations is an ubiquitous phenomenon in neural systems and may represent a key neuronal mechanism underlying cognitive or perceptual processing (Buzsáki 2006). Neuronal oscillations are traditionally measured by EEG, recordings of local field potentials (LFP), or multi-unit recordings. Beyond the depiction of this neuronal synchronization, identifying driver-response relationships between interconnected brain sources and understanding their directed interactions and dynamics can also inform the functional architecture of sensory and cognitive processing, in both healthy and diseased brains (Bressler 1995). There are various measures that have been developed to identify driven and driving interactions between brain sources. These measures vary from linear to nonlinear, bivariate to multivariate, and many rely on the Granger causality principle (Granger 1969). This approach quantifies improvement in the predictions of a time series, given its past, when information from the past of another time series is considered (Baccalá & Sameshima 2001; Kamiński & Blinowska 1991). These measures, which are based upon statistical dependencies in data over time, are thought to provide measures of directed functional connectivity. Another approach, DCM (David and Friston 2003), is used to infer (directed) effective connectivity, that is, how one source or neural system influences another. The main distinction between DCM (model-based) and Granger-based (data-based) methods is that DCM is based on biologically plausible neural mass models that are inherently causal in nature. In other words, the question is not whether there is (Granger) causality – but which (causal) models best accounts for data. This enables one to identify how a system of pre-specified neuronal populations generates the measured signal (Schoffelen and Gross 2009), and to compare different hypotheses or architectures in terms of their model evidence.

Measuring connectivity at the scalp level can be informative but one has to be careful about its interpretation in terms of brain dynamics. This is because scalp data sees neuronal sources through a specific ‘lens’ which distorts, mixes and loses information about the exact location of the underlying sources. A fundamental problem with scalp recordings is electrical conduction through the head volume. This means that instead of recording brain activity from one specific brain source, each sensor measures a linear superposition of signals from all over the brain. This mixing introduces instantaneous correlations in sensor data, so that the interpretation of directed connectivity has to proceed with caution because spurious connectivity patterns can arise. In short, scalp recordings provide an indirect measure of source activity (with rather low signal to noise ratio), which is not easily interpretable. A critical assessment of directed connectivity measures based on EEG recordings can been found in Haufe et al. (2013). The authors report a series of simulations to assess the sensitivity of sensor-based functional connectivity when inferring source interactions from synthetic EEG recordings.

To make inferences about directed connectivity among brain sources one can either apply source reconstruction techniques to estimate source activity or use intracranial EEG (iEEG) data from electrodes implanted in human subjects (e.g., patients with brain tumours and epilepsy). Invasive iEEG recordings are difficult to obtain but they have been of great help, not only as a part of pre-surgical evaluation for patients, but also in the study of responses induced by cognitive tasks. These responses would be almost impossible to study with high precision on the scalp level. Finally, invasive (but rare) electrophysiological recordings can be used to validate the reconstruction and modelling of (readily available) sensor level data. This is one of the aims of our paper.

There are two prevalent approaches to measuring directed connectivity in the spectral domain. These are exemplified by (data-based) DTF (Kamiński and Blinowska 1991) and (model-based) biologically informed DCM (Friston et al. 2003). The first approach generalizes the concept of Granger causality to the spectral domain. It has been applied to iEEG recorded from patients with epilepsy: i) around the seizure onset, to identify the putative epileptogenic zone (van Mierlo et al. 2011; Papadopoulou et al. 2012) or ii) during the performance of cognitive tasks, to investigate distributed neuronal processing (Brázdil et al. 2009; Flinker et al. 2015). DTF has also been used to infer directed functional connectivity between reconstructed sources (RS) in humans (Dai et al. 2012) and from intracranial recordings of monkeys (Liang et al. 2000). Similar approaches have addressed connectivity at the source level using Independent Component Analysis (Haufe et al. 2010), where DTFs have also been computed (Gómez-Herrero et al. 2008; Cantero et al. 2009). In contrast to these data-based measures, DCM uses neurobiologically plausible models that are fitted to empirical observations, which are then subjected to Bayesian model comparison or selection (BMS). BMS allows one to evaluate competing hypotheses (or architectures) in terms of their Bayesian model evidence or marginal likelihood. In brief, DCM treats the brain as a nonlinear dynamical system that receives inputs and generates outputs. In this setting, an experiment is regarded as a perturbation (induced by the inputs) of coupled electromagnetic sources, which produces source-specific responses (Kiebel et al. 2009). The basic idea behind the method is to model the influence of each source on others – and identify the mechanisms that underlie distributed network responses. Dynamic causal modelling has been applied to both functional magnetic resonance imaging (fMRI) and magnetoencephalography (MEG)/EEG data.

DCM for MEG/EEG is based on a spatiotemporal generative model of electromagnetic brain activity, where the temporal dynamics are described by neural mass models of equivalent current dipole (ECD) sources, and their spatial expression at the sensor level is modelled by parameterized lead-field functions. Generally a DCM comprises a model of interacting cortical sources, where each source corresponds to a canonical circuit of neural populations, and its electromagnetic output is generated by the modeled average depolarization of pyramidal cell populations. These electromagnetic outputs are then passed through an electromagnetic model of the head, accounting for volume conduction effects, to finally generate predictions at the M/EEG sensor level (Fastenrath et al. 2009). This process is called the forward problem, as opposite to the inverse problem which infers the activity in the brain starting from scalp recordings. Equipping neuronal models with a lead field effectively subsumes the source reconstruction problem into model inversion or fitting. In other words, DCM can estimate directed effective connectivity among sources using sensor data directly. DCM has been extensively applied to sensor space data to infer directed effective connectivity in healthy and diseased subjects (e.g., Garrido et al. 2008; Garrido et al. 2009; Herz et al. 2012; Herz et al. 2013; Herz et al. 2014). It has also been applied to LFP recordings in rodents (Moran et al. 2009; Moran et al. 2011; Moran et al. 2015) and intracranial electroencephalographic (iEEG) in humans (Papadopoulou et al. 2015). In some applications, DCM is applied to source reconstructed data in source space, as opposed to modelling responses in sensor space. This allows one to make inferences about connectivity among a predefined set of sources, without having to consider all the sources generating sensor data (e.g. Boly et al. 2012). This is the approach we adopt in the current paper, as we wanted to focus on a subset of sources for which we had invasive or direct recordings.

In this work we analyzed ECoG and source reconstructed data from one monkey during wakefulness and propofol anaesthesia. Our aims were twofold; first, we wanted to see whether directed connectivity in the frequency domain between cortical sources reconstructed from scalp EEG is qualitatively similar to estimates based on ECoG recordings, using both DTF and DCM. Our second focus was on how the information flow between two pre-specified sources (frontal and parietal) was modulated in wakefulness and sedation.

It is worth mentioning that our aim is to compare the connectivity results obtained by reconstructed sources on one hand and the corresponding intracranial recordings on the other; a comparison of data-driven (DTF) and biophysical (DCM) models for directed dynamical connectivity is not the scope of the present work.

## Methods

### Data

These data are part of a dataset collected at a workshop titled “Controversies in EEG source imaging”, held in August 2014 at the University of Electronic Science and Technology in Chengdu, China, with the aim of discussing the major issues at stake when brain activity is recorded or modelled as electrical potentials. All the simulations and data are available from the following website http://neuroinformation.incf.org/ and will be described in detail in a technical report. Specifically for this study we used publicly available data (http://neurotycho.org/) that were originally analyzed and published in Yanagawa et al. (2013). ECoG and EEG signals were simultaneously recorded from the same monkey (*Macaca Mulatta*). The monkey was implanted with a 128 channel ECoG array that covered the lateral cortical surface of the left hemisphere with 5 millimeter spacing. EEG signals were recorded from 19 channels. The EEG electrodes locations conformed to the 10-20 system without Cz (to avoid interference with an ECoG connector). ECoG and EEG data were sampled at 1000Hz. The monkey was seated in a primate chair with eyes closed and both arms constrained – and injected with an anaesthetic drug (propofol) during the recording to induce loss of consciousness.

In the following we report the steps for the leadfield reconstruction. Using BrainSuite2, a T1 MRI was corrected for intensity bias and segmented into tissues (i.e. grey and white matter) and cerebrospinal fluid. The white/grey matter interface was chosen as the source space model for EEG/ECoG, i.e., each node of the mesh was a potential source. The head was then divided into brain (enclosed by the pial surface), brain plus surrounding cerebrospinal fluid, skull and skin. This segmentation was checked and adjusted manually by an expert. The volume conductor model was based on the above segmentation, assuming constant electrical conductivities within each compartment. The skull-to-other conductivity ratio was set to 1/25. 1mm-thick silicone strips (housing the ECoG electrodes) were also included in the model because silicone has very low conductivity and can influence EEG signals. An X-ray 2D image was spatially registered to the pial surface. The transformed electrode positions were then projected onto the 3D pial surface. The silicone stripes were modelled according to Figure 1 in Nagasaka et al. (2011). These were modelled as a grid of 1-mm thick silicone rings of 3.5 mm radius, each surrounding an electrode of 2.0 mm radius. The conductivity of the silicone was set to a negligible value relative to the other compartments. The EEG electrodes were manually located on the monkey’s scalp using IMAGIC (<u>www.neuronicsa.com</u>) and projected onto their corresponding mesh faces.

**Figure 1.**
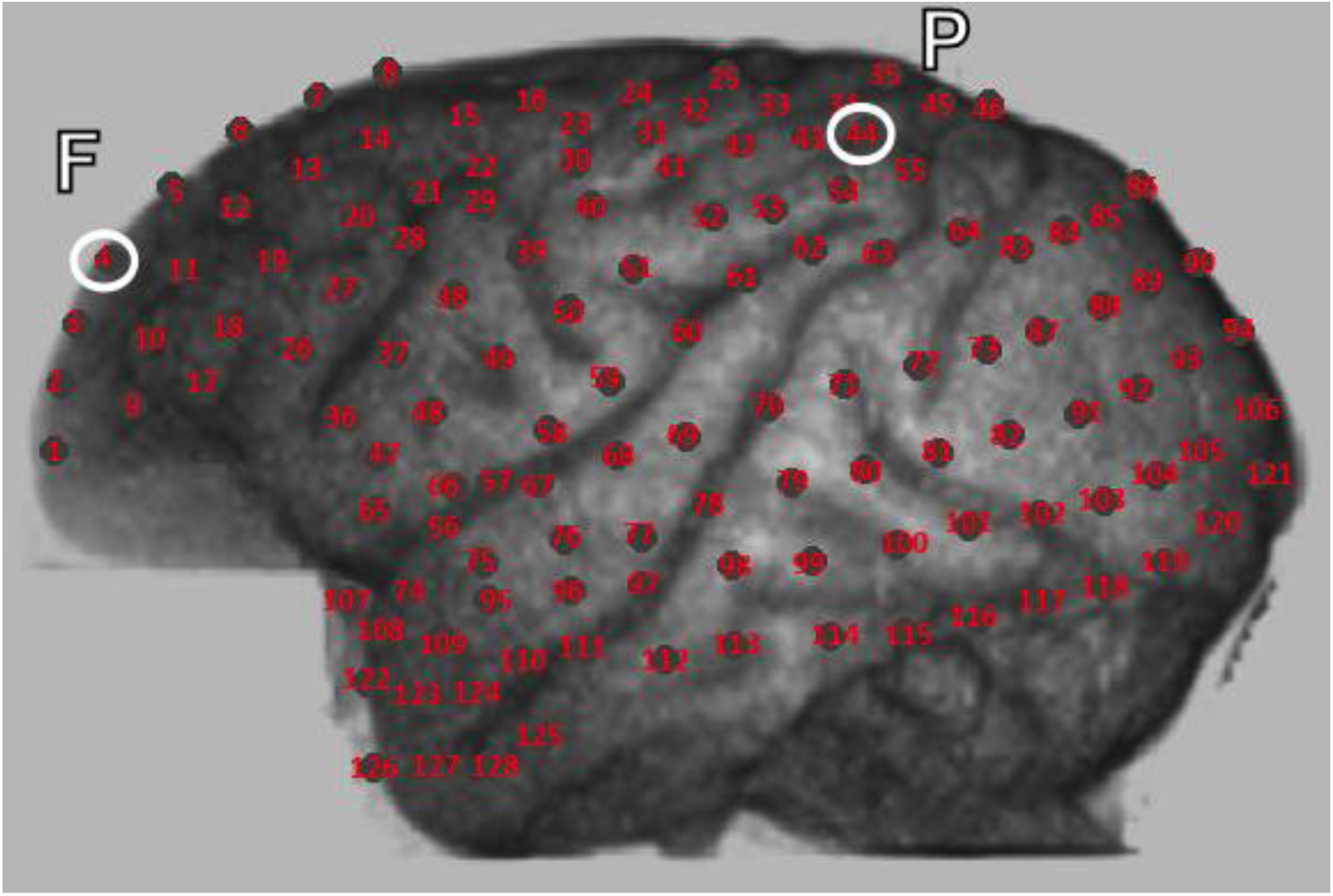
Layout of the ECoG contact locations. The frontal (F) and parietal (P) channels used in this study are indicated by white circles.

Tetrahedral meshes were created from the surfaces of the head model using Tetgen 2.0 (open source). Both EEG and ECoG lead fields were calculated using NeuroFEM, a program for computing lead fields using the Finite Element Method, which is part of the SimBio software package (*SimBio Development Group. “SimBio: A generic environment for bio-numerical simulations”, <u>https://www.mrt.uni-jena.de/simbio</u>*). Source reconstruction in the time domain (for the EEG data) was performed by LORETA (free academic software for source localization of EEG data: http://www.uzh.ch/keyinst/loreta)(Pascual-Marqui et al. 1994). The estimated current sources were constrained to be perpendicular to the cortical surface. No absolute value or norm was taken for the dipole or the resulting data, so no period doubling effects are to be expected. EEG sources were reconstructed in both hemispheres. For this study we only retained the RS nearest to the ECoG channels considered in the connectivity analyses. The correspondence between cortical and reconstructed activity was assessed by means of canonical correlation analysis to provide a goodness of fit measure (results not shown here).

The pre-processing steps for both ECoG and RS included average reference removal, notch filtering at 50Hz, artefact removal by visual inspection and local detrending with the L1 norm technique (Kim et al. 2009). In the current validation study we restrict our analysis to a single pair of sources, a frontal source (**F**) and a parietal source (**P**), as indicated in Figure 1. This choice was motivated by a previous study using RS from scalp EEG recordings in humans that measured directed connectivity between cortical sources in these areas (Boly et al. 2012), and functional connectivity in anesthetized macaque monkeys (Moeller et al. 2009; Barttfeld et al. 2015). 30 seconds of brain activity were used for each condition (wakefulness and anaesthesia).

The spectra of the two channels in the two conditions are reported in Figure 2, together with the spectra of the data modelled with an autoregressive model of the composite system of the two sources, of order 7, as the one used for DTF.

**Figure 2.**
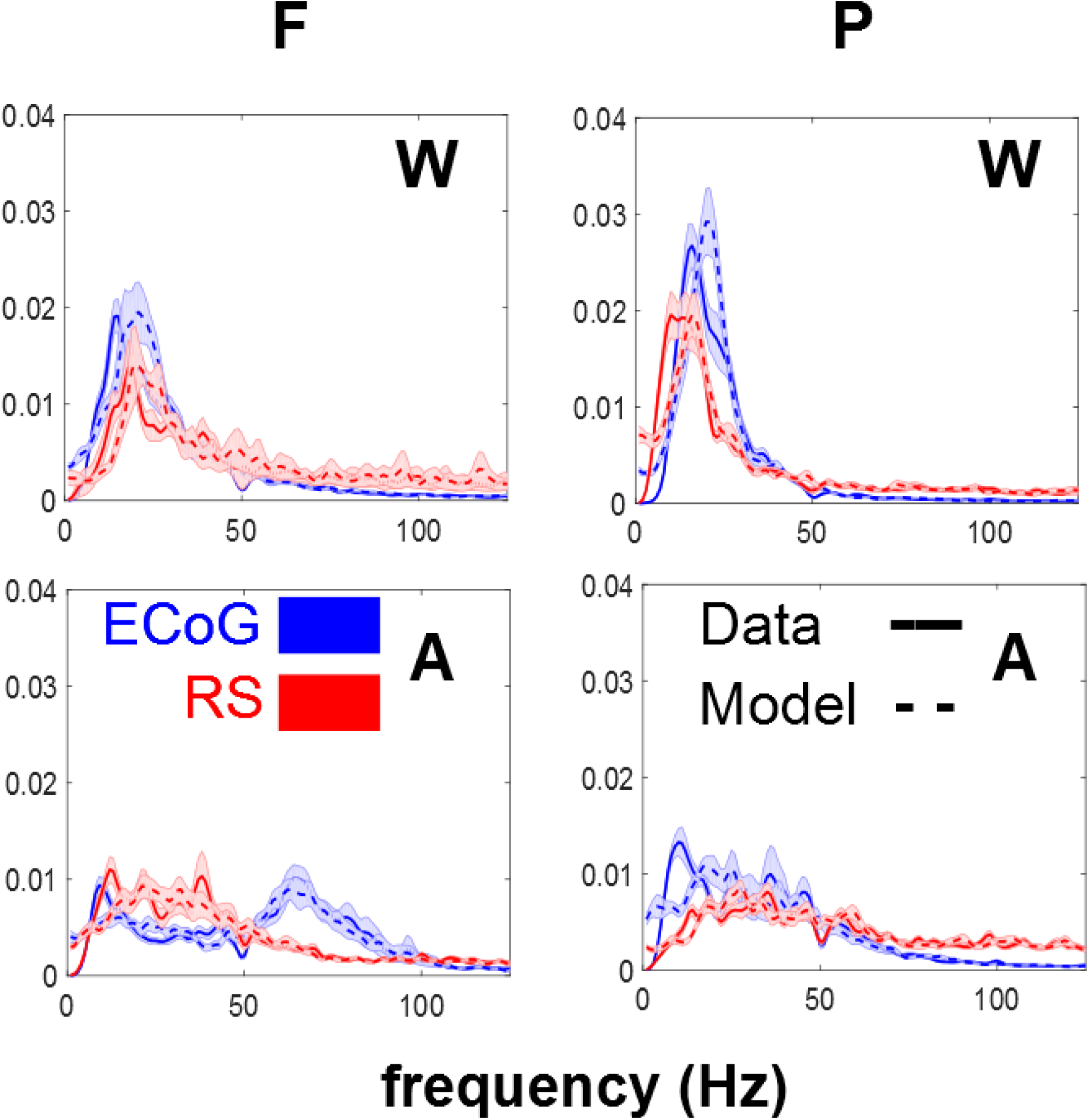
Power spectral densities of real data (full line) and data simulated with the coefficient of an autoregressive model of order 7 of the real data (dashed line) for ECoG (blue) and reconstructed sources (red). Left column: frontal source (F). Right column: parietal source (P). Top panels: wakefulness (W). Bottom panels: anaesthesia (A).

As shown in <u>http://wiki.neurotycho.org/EEG-ECoG_recording</u> EEG signals don’t include high frequency (> 60Hz) components of the ECoG signal.

### Directed Transfer Function

The DTF is a multivariate directed functional connectivity measure, usually based on an autoregressive model (AR) in the frequency domain (Kamiński and Blinowska 1991). The AR model is of the form

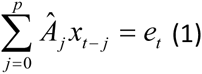

where *x*_*t*_ =(*x*_1,*t*_, *x*_2,*t*_,…., *x*_*k*,*t*_) is a vector of *k*-channel multivariate processes, *e*_*t*_ = (*e*_1,*t*_,*e*_2,*t*_,…‥,*e*_*k*,*t*_) is a vector of multivariate uncorrelated white innovations or noise processes and 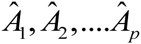 are the *k* × *k* matrices of model coefficients. Multiplying both sides of (1) by 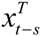 and taking the expectation returns the coefficients 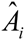, as follows

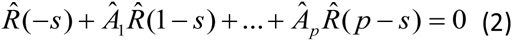

where 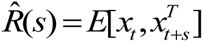 is the covariance matrix for time lag *s*. To characterize Granger causal coupling between signals in the spectral domain, the Fourier transformation of equation (1) is calculated, where the transform functions are of the form 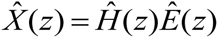 where 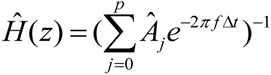 The DTF then is derived from the transfer matrix and can be expressed as:

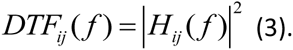

Usually, the DTF is normalized with respect to the incoming to the incoming information flow so that it takes the form

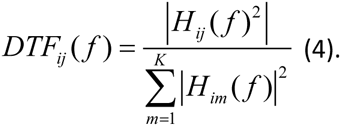

Consequently, the element *H*_*ij*_ (*f*) of the matrix describes the connection between the *j*-th input and the *i*-th output at each frequency. The values of the normalized DTF are located in the range [0, 1] where a high value indicates a greater information transfer in the direction *j* → *i* and a low value indicates little or no transfer. For the present study we used 7 as the autoregressive model order, as determined by the Bayesian Information Criterion.

In a recent Opinion paper, Kaminski and Blinowska (2014), the inventors of DTF, postulated that this measure is not sensitive to volume conduction, since it is insensitive to phase shifts. However, while it is true that a phase shift in sensor data indicates information transfer, no inference can be made about where the implicit sources are located, except in special cases in which the experimental protocol or the anatomy ensure that the activity of a single source is expressed at a single sensor (Plomp et al. 2014).

As mentioned above, DTF is applied to two sources, a frontal and a posterior one, for each level of consciousness, using 15 non-overlapping segments of 2 seconds.

### Dynamic Causal Modelling

For this study we used DCM for cross-spectral density (CSD), which is a generalization of DCM for steady state responses. All our analyses used the standard procedures described in (Friston et al. 2012). CSD is the Fourier transform of the cross-correlation function and can be thought of as reporting the correlations at each frequency. CSD therefore describes the similarity between two signals, that is, how much power is shared for each frequency.

The neural mass model used here was the LFP variant. This particular neural mass model has been used previously in modelling intracortical local field potentials from rats, to assess changes in directed effective connectivity under pharmacological manipulations (Moran et al. 2009; Moran et al. 2015). It has also been used as a generative model for non-invasive EEG studies, in source-reconstructed data from frontal and parietal cortices during normal wakefulness, propofol-induced mild sedation and loss of consciousness in humans (Boly et al. 2012).

One can regard each neural mass as a cortical source, where each source comprises three subpopulations that contribute to the ongoing dynamics. These subpopulations include spiny stellate cells in the granular layer and pyramidal cells and inhibitory interneurons in supragranular layers.

Each of the subpopulations is modelled with pairs of first order differential equations of the following form:

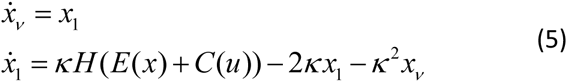

The column vectors *x*_*v*_ and *x*_1_, correspond to the mean voltages and currents where *E*(*x*) and *C*(*u*) correspond to endogenous and exogenous inputs respectively that the presynaptic input to each subpopulation comprises (see Moran et al., 2009).

The nodes (sources) of DCM model sources in the brain are connected by (extrinsic) forward and backward connections according to anatomical connectivity rules established in (Felleman and Van Essen 1991). Feedforward connections target the granular layer, while feedback connections target the superficial and deep layers (Bastos et al. 2012). More details about the different models that can be used within the DCM framework can be found in (Moran et al. 2013).

Here, we first use DCM to test hypotheses about the connectivity architecture between the two sources of interest in frontal and parietal regions. We tested two physiologically plausible models. Our first model connects the parietal to the frontal source by forward connections and frontal to parietal with backward connections, while the second model constitutes the reverse architecture (Figure 3).

**Figure 3.**
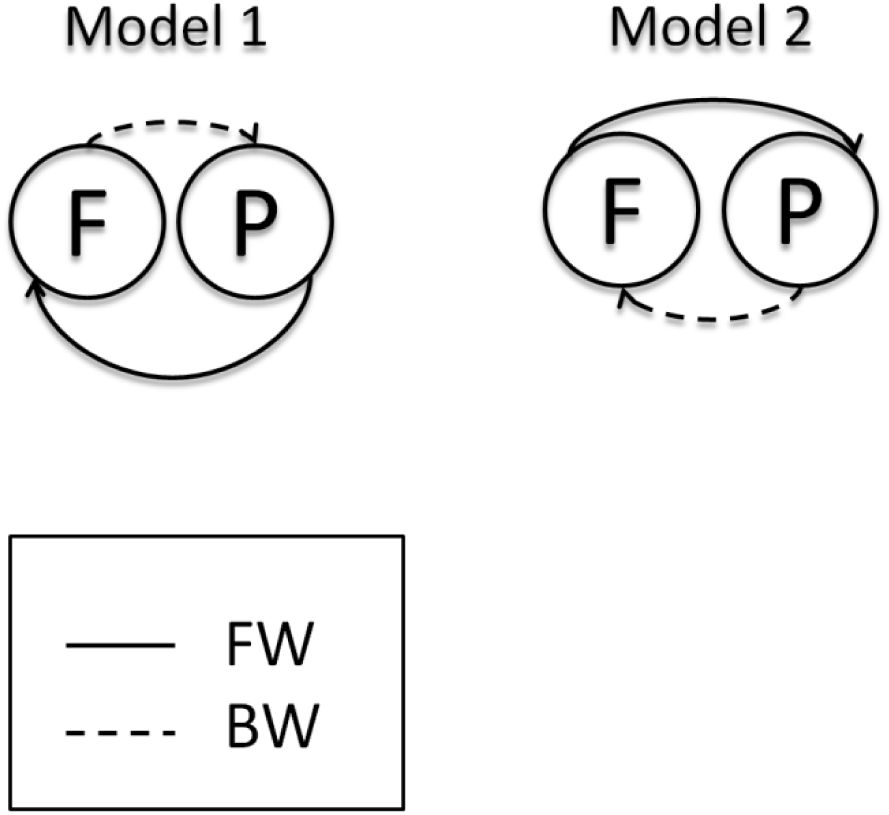
The two architectures for connections between the sources of interest tested with DCM.

The designation of fronto-parietal and parieto-frontal connections as backward and forward is based on the functional asymmetries in the anatomy and physiology of projections – extrapolating from the visual system. A brief review of this evidence, from the point of view of the extended motor system can be found in (Shipp et al. 2013).

We inverted the two models using both sets of empirical data and then performed (fixed effects) Bayesian model selection (BMS) to identify the most likely model. We then modelled the condition-specific effects under the best model, corresponding to wakefulness and anaesthesia. These effects were modelled in terms of changes in intrinsic and extrinsic connections relative to the first condition (wakefulness) (Figure 4).

**Figure 4.**
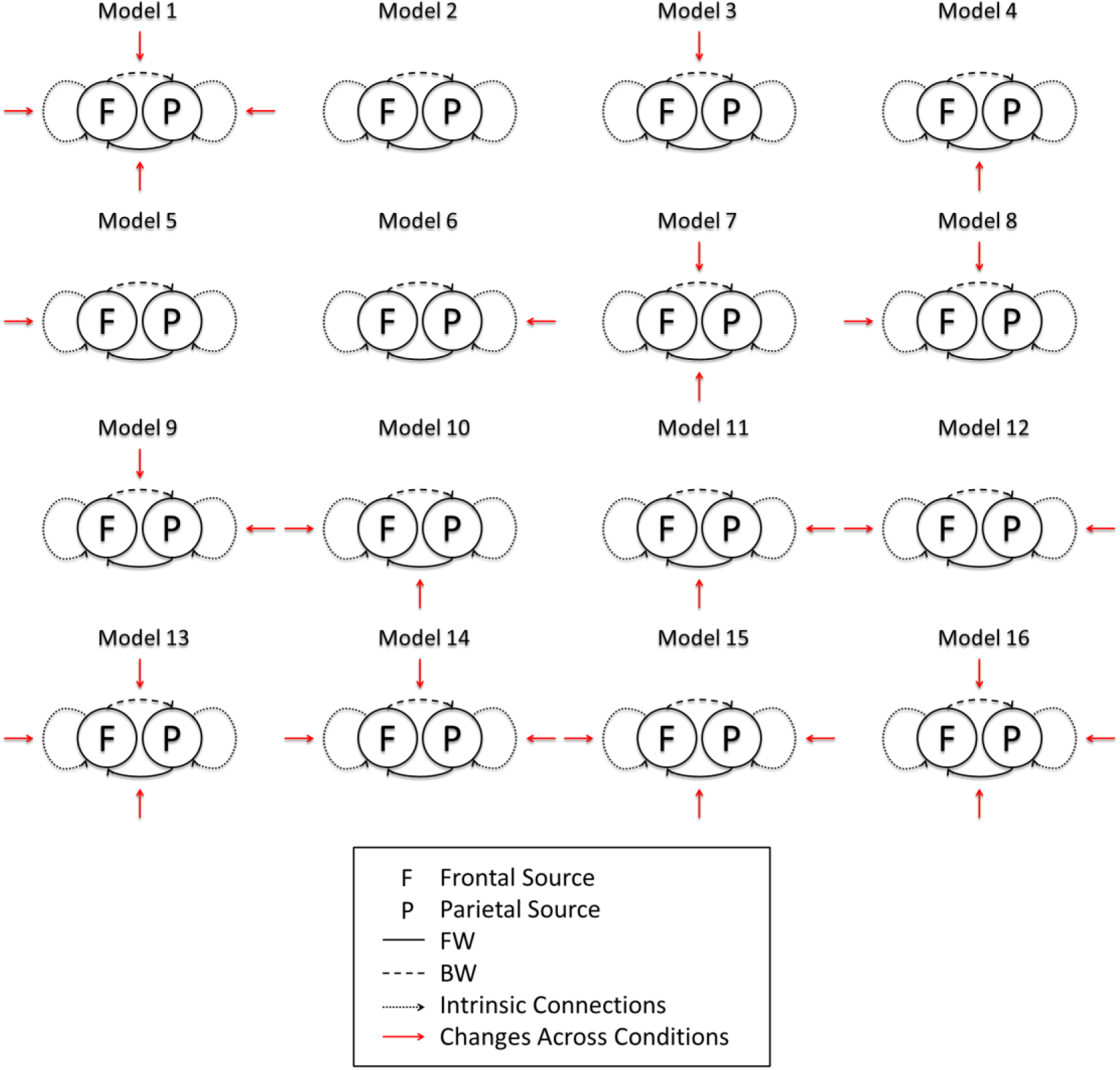
The sixteen possible models tested by DCM to explain changes in connectivity from wakefulness to anaesthesia.

## Results

In this study we evaluated directed connectivity in the frequency domain between two sources located in frontal and posterior brain regions, and determined how the information flow between the two sources is modulated by anaesthesia. This evaluation used both ECoG and reconstructed source activity, enabling us to assess the validity of connectivity estimates based upon non-invasive EEG signals.

DTF quantifies information flow across brain areas for each frequency bin. The curves for each condition and modality are reported in (Figure 5). We have assessed the significance of the modulations corresponding to the spectral interval [3 40] Hz with a nonparametric Wilcoxon Rank Sum test. Significant decrease during loss of consciousness is reported in the connectivity from the parietal to the frontal source, for both the ECoG and reconstructed EEG source activity (P<0.02, FDR corrected). The other modulations, tested across consciousness state and across imaging modalities, were not significant.

**Figure 5.**
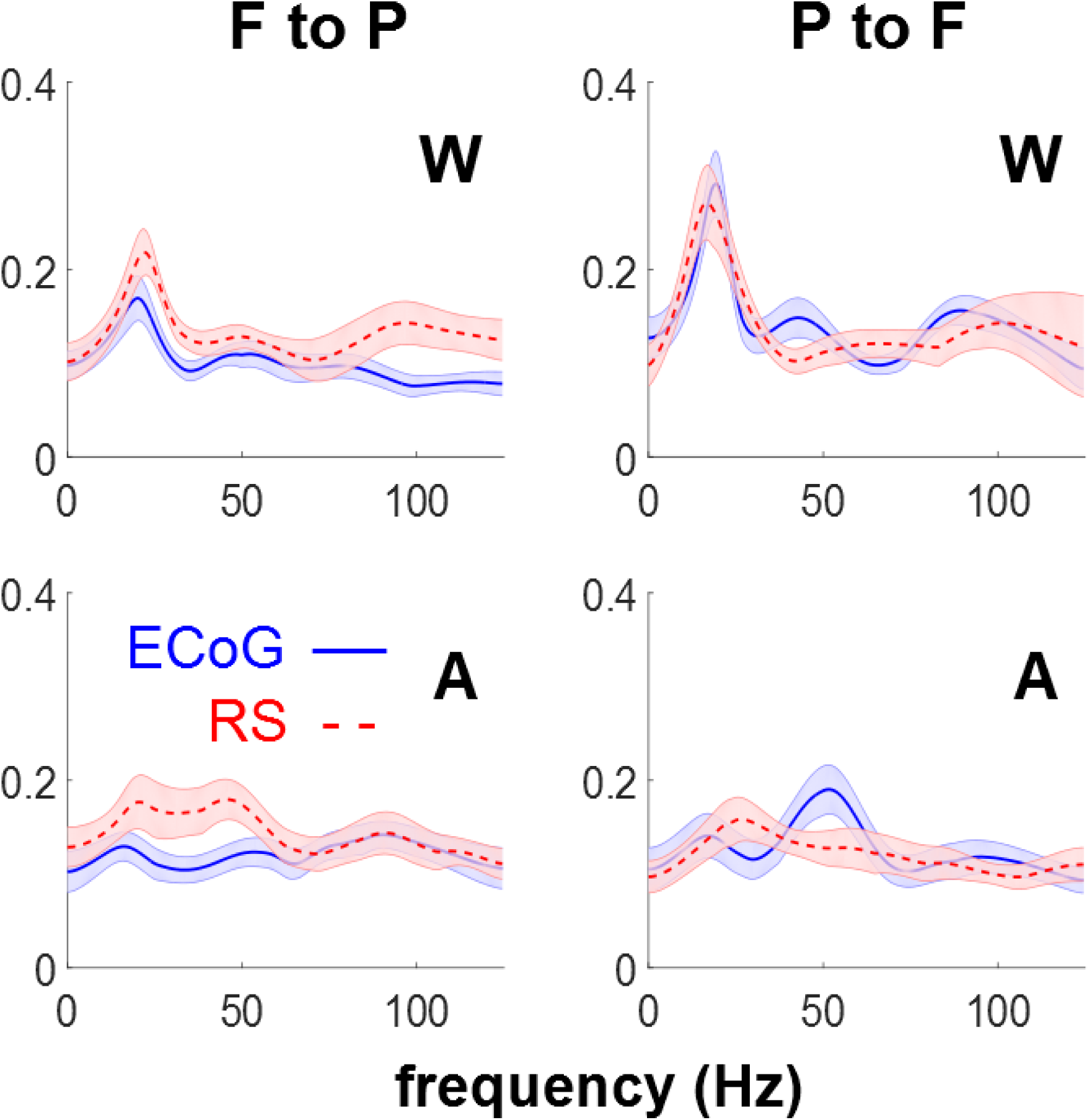
DTF plotted against frequency in the two directions in Wakefulness (W) and Anaesthesia (A) for ECoG and RS. Shaded areas indicate standard errors.

DCM and BMS of the directed effective connectivity between the same sources identified model 1 as the most plausible, with a forward connection from the parietal to frontal region and backward connections from the frontal to the parietal region (Figure 3). The difference between the best and next best model was much greater than three reflecting strong evidence in favour of the first model over competing hypotheses. The same winning model was identified for ECoG and reconstructed EEG source activity.

For the second part of our DCM analysis, we modelled condition-specific effects in terms of all the possible combinations of condition-specific changes in the forward connections, the backward connections, neither or both. BMS identified model 1 as the winning model (Figure 4). This model allows for changes all the connections. As before, the same winning model was identified for both ECoG and reconstructed EEG sources. The differences in log evidence among the four models were comparable for the invasive data and to the reconstructed EEG data. This suggests that there is roughly the same amount of information in both modalities when it comes to disambiguate the models or hypotheses. This is reflected also the posterior probabilities (left panel of Figure 6) over models, which are also comparable.

**Figure 6.**
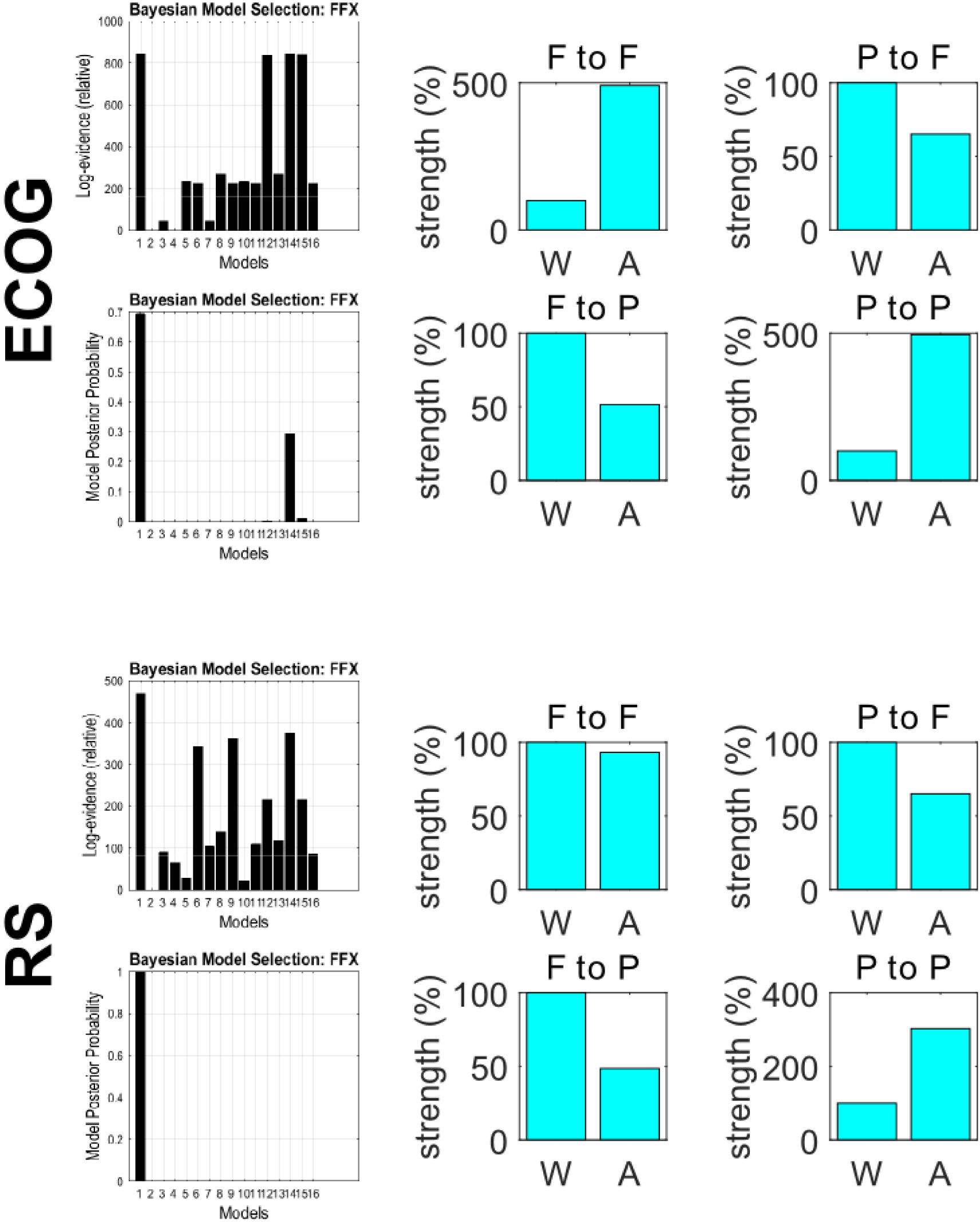
Log evidences and posterior probabilities (left) and changes in connectivity in the winning model (model 1, right) for ECoG sources (top) and reconstructed sources (bottom) across the two conditions: Wakefulness (W) and Anaesthesia (A), as estimated by DCM.

Looking at the condition-specific effects on the extrinsic connectivity (Figure 6, right panel), the parameter estimates based upon the ECoG data concur with the changes in DTF; namely, a decrease is seen in both forward connectivity from the parietal to the frontal source, and in backward connectivity from the frontal to the parietal source. At the same time a strong increase in self connections in anaesthesia is reported in both sources for ECoG; a slight decrease in the frontal source and a moderate increase in the parietal source for reconstructed activity. These changes are relative to the 100% connectivity strength in the wakefulness condition.

One interesting aspect of DCM is that we can estimate the DTF implicitly from the condition-specific effects on the parameters. In other words, given the model parameters, we can compute the associated directed transfer functions between the sources, as shown in Figure 7. This figure uses the same format as Figure 5. The fact that directed transfer functions (and Granger causality) can be derived from the DCM results speaks to the fact that Granger causality and directed transfer functions are essentially data features (hence data-led measures), and not the model attributes responsible for directed information flow. It is pleasing to note that, qualitatively, the data-based DTFs and those based upon DCM parameter estimates show the same dependency on experimental condition. Higher DTF values at frequencies higher than the main peak are observed in from the frontal to the parietal source for electrocorticogram but not for the reconstructed sources. The forms of the DTFs are more constrained under DCM, because they have to be produced by a biologically plausible mechanism. Furthermore, the DCM transfer functions have been modulated by the spectral power of the innovations (which is also estimated). Note that the autoregressive evaluations of DTFs do not estimate the spectral density of the innovations, which are assumed to be white (see equation 1).

**Figure 7.**
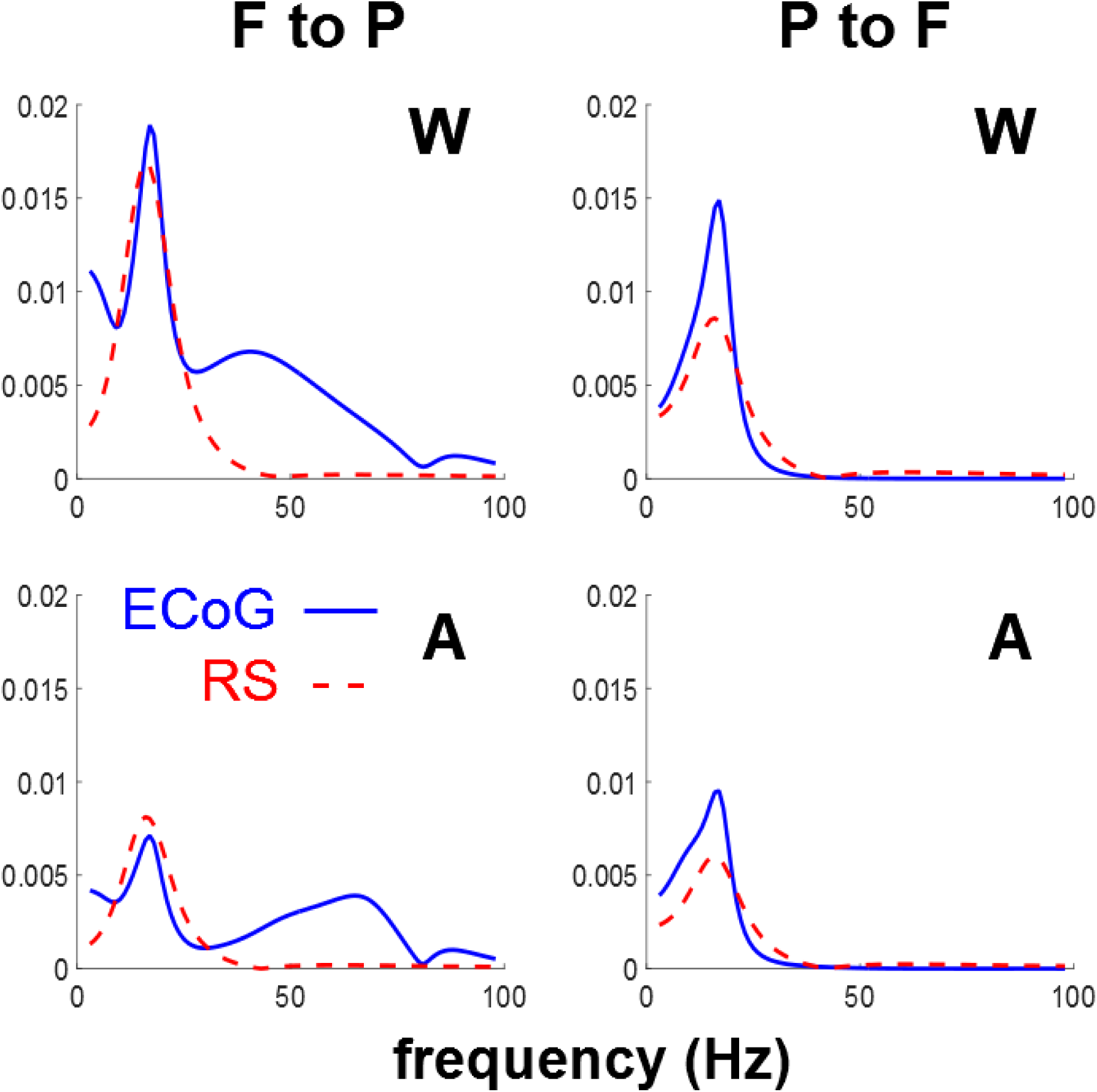
Directed Transfer functions obtained from DCM under the winning model in the two conditions: Wakefulness (W) and Anaesthesia (A) for ECoG and RS.

## Discussion

In a previous study that analyzed these data with directed functional connectivity, all possible pairs of ECoG sources (with a bipolar montage) were considered (Yanagawa et al. 2013). Functional connectivity differed significantly between conscious and unconscious states in all combinations of cortical sources, with the most dramatic change occurring for the transfer functions that fell into a specific spectral domain across conditions. This motivated the authors to look for large-scale inter-region interactions over the entire cortex by grouping the bipolar channels in 8 cortical regions, after which spectral Granger causality was computed for each pairwise combination. The changes in connectivity patterns after this grouping confirmed that the spectral changes due to modulations of consciousness affected large-scale communications across the entire cortex.

Here we focused on a pair of sources since in many experiments, in particular event-related ones, only a few sources are considered, and in order to apply DCM between two regions known to play a distinct joint role in wakefulness versus anaesthesia.

In this study, we have shown that the directed connectivity in the frequency domain between cortical sources reconstructed from scalp EEG is qualitatively similar to, and statistically undistinguishable from, the connectivity inferred directly from cortical recordings. The modulations of DTFs across frequency are qualitatively the same (although in a few cases the peaks differ slightly in position or width). Concerning the effects of the anaesthesia, the same pattern emerged from electrocorticographic and reconstructed sources, with a decrease in the information flow from the parietal to the frontal source. This modulation is in general agreement with previous literature (Lee et al. 2009; Ku et al. 2011; Boly et al. 2012). This comparison must stay qualitative since the studies mentioned above consider human subjects and scalp EEG.

Dynamic causal modelling produced Bayesian model comparisons that were consistent between electrocorticograms and reconstructed sources. These models explained the decrease in coupling from parietal to frontal sources in terms of condition-specific changes in extrinsic (forward and backward) connectivity with the frontal source as well to changes in intrinsic connectivity at both sources. In these analyses, Bayesian model selection based on the invasive and non-invasive data was again consistent; however, quantitative connectivity changes following inversion of the EEG and ECoG data showed opposite changes in extrinsic and intrinsic connectivity but similar directed transfer functions. We mention this discrepancy to illustrate that the quantitative estimates of effective connectivity can, in some instances, depend upon the nature of the data, especially when there is a conditional dependency among parameter estimates. In principle, one would base their inferences on all the data at hand and model both the ECoG and EEG data together. In this setting, the most precise or informative data would supervene in terms of model comparison and parameter estimates (the model comparison results in Figure 5 would suggest that the ECoG data were more precise). In more realistic DCM analyses, one generally includes several sources to disambiguate between explanations based upon reciprocal changes in intrinsic and extrinsic connectivity. One of the characteristics of DCM is that it can also model hidden sources; for example the thalamic sources in Boly et al. (2012). The inclusion of hidden sources is sometimes required to adjudicate among different hypothetical architectures, using Bayesian model selection in the usual way. Crucially, this is not an option with data-led measures of directed functional connectivity. Further discussion of the relationship between data-driven functional connectivity in the spectral domain and DCM based measures of effective connectivity can be found in Friston et al. (2014).

The adequacy or quality of any model is generally established through Bayesian model comparison. Good models have a high evidence and entail a level of complexity that is suitable for the data at hand. The DCM of source activity has been refined over many years and provides the appropriate level of detail – in terms of the number of sources and parameters. These parameters include not just aspects of the underlying neuronal (connectivity and synaptic) architecture but how neuronal activity is measured. For example, the contribution of different neuronal populations to different sorts of sensors is accommodated through free parameters, that scale the relative contributions (with a prior bias towards superficial populations).

Being aware of the limitations of single-subject studies, we do not infer any pathophysiological explanations from our results. Also, a task protocol with more localized sources would definitely provide additional insight. Nonetheless this unprecedented recording setup provides a valid support for the exploratory analysis that we performed with the sole protocol available at the moment, which allowed us to explore modulations in steady-state activity. It is worth to recall that whenever activity has to be estimated or disambiguated with a fine spatial resolution, a large number of scalp electrodes is recommended.

Our provisional results suggest that directed connectivity in the frequency domain between cortical sources reconstructed from scalp EEG is qualitatively similar to the connectivity inferred directly from cortical recordings, using both functional and effective connectivity measures. These findings advocate that inferences about directed connectivity based upon non-invasive electrophysiological data can have construct validity in relation to invasive constructs.

## Acknowledgements

The leadfield reconstruction was performed by Pedro A. Valdés Hernandez (Cuban Neuroscience Center). For the source reconstruction and the processing of the data used in this paper we wish to thank also Pedro Valdés Sosa, Jorge Bosch-Bayard, Esin Karahan and Peng Ren. We also acknowledge the assistance of Gregor Strobbe in the early phase of this study. We wish to thank Pedro Valdés Sosa for the idea of setting up a workshop on EEG source imaging and a special issue connected to it, and for coordinating the effort of collecting and processing the datasets KJF is funded by a Wellcome Trust Principal Research Fellowship (Ref: 088130/Z/09/Z)

